# Clinical metagenomics bioinformatics pipeline for the identification of hospital-acquired pneumonia pathogens antibiotic resistance genes from bronchoalveolar lavage samples

**DOI:** 10.1101/2020.02.26.966309

**Authors:** Maud Tournoud, Etienne Ruppé, Guillaume Perrin, Stéphane Schicklin, Ghislaine Guigon, Pierre Mahé, Vladimir Lazarevic, Sébastien Hauser, Caroline Mirande, Albrice Levrat, Karen Louis, Gaspard Gervasi, Jacques Schrenzel

## Abstract

**Background:** Shortening the time-to-result for pathogen detection and identification and antibiotic susceptibility testing for patients with Hospital-Acquired and Ventilator-Associated pneumonia (HAP-VAP) is of great interest. For this purpose, clinical metagenomics is a promising non-hypothesis driven alternative to traditional culture-based solutions: when mature, it would allow direct sequencing all microbial genomes present in a BronchoAlveolar Lavage (BAL) sample with the purpose of simultaneously identifying pathogens and Antibiotic Resistance Genes (ARG). In this study, we describe a new bioinformatics method to detect pathogens and their ARG with good accuracy, both in mono- and polymicrobial samples.

**Methods:** The standard approach (hereafter called TBo), that consists in taxonomic binning of metagenomic reads followed by an assembly step, suffers from lack of sensitivity for ARG detection. Thus, we propose a new bioinformatics approach (called TBwDM) with both models and databases optimized for HAP-VAP, that performs reads mapping against ARG reference database in parallel to taxonomic binning, and joint reads assembly.

**Results:** In in-silico simulated monomicrobial samples, the recall for ARG detection increased from 51% with TBo to 97.3% with TBwDM; in simulated polymicrobial infections, it increased from 41.8% to 82%. In real sequenced BAL samples (mono and polymicrobial), detected pathogens were also confirmed by traditional culture approaches. Moreover, both recall and precision for ARG detection were higher with TBwDM than with TBo (35 points difference for recall, and 7 points difference for precision).

**Conclusions:** We present a new bioinformatics pipeline to identify pathogens and ARG in BAL samples from patients with HAP-VAP, with higher sensitivity for ARG recovery than standard approaches and the ability to link ARG to their host pathogens.

## Background

Over the past decade, methods for capturing and analyzing all DNA present in a sample have become available, opening the metagenomics era. Sifting through the reads from eukaryotic, viral and bacterial DNA is now conventional, e.g. SURPI or One Codex platforms [1, 2], whereas obtaining separate genome assemblies, and even better functional analysis including resistance to antibiotics for each detected pathogen, remains a challenge in the context of hospital-acquired and ventilator-associated pneumonia (HAP-VAP). Obtaining antimicrobial susceptibility testing results earlier than the minimum 6-8 hours from the culture is a prerequisite to allow an earlier adjustment of antibiotic therapy and possibly better patient outcome [3] [4]. This would contribute to prevent multi-drug resistant bacteria from dissemination [5] and to limit patient mortality caused by inadequate therapy [6].

Metagenomic analysis methods for taxonomic analysis are continually improving and fall into two broad groups: read-based and assembly-based analyses. In the read-based group, a first approach consists in taxonomic read binning using kmers (e.g. Kraken [7]) or Ferragina-Manzini index (e.g. Bowtie2 [8], Centrifuge [9]), followed by taxonomic analysis based on lowest common ancestor. The integrated MEGAN [10] pipeline implements these two steps. Alternatively, it is possible to perform taxonomic profiling using marker genes (e.g. MetaPhlAn2 [11], TIPP [12]).

For the assembly-based methods, the CAMI challenge [13] has identified MEGAHIT [14] as one of the best choices, among many alternatives, to obtain a metagenome assembly. Then, meta-assembly contigs can be assigned to species using similarity-based approaches or evaluating their nucleotide composition and abundance patterns [15, 16]. Expert supervision remains, however, still required to detect mis-assemblies [17], and despite their great performance, the computational burden of these methods is still demanding.

In this article, we want to go beyond taxonomic analysis and present a bioinformatic method able to identify pathogens involved in HAP-VAP infections, detect the presence of Antibiotic Resistance Genes (ARG) and link them to their host bacteria. In particular, we focus on the case of ARG which can suffer from low detection performance, because they are either absent from the genome reference database or present in several genomes. Moreover, assuming that an ARG is present in the metagenome, it is important from a clinical perspective to distinguish between its presence in commensal bacteria and the more clinically relevant situation where the ARG is harbored by one or several species of pathogenic bacteria. This objective is also addressed by MetaCherchant [18] which aims at extracting the genomic environment of ARG detected in a metagenome and thus infer the link with the host bacteria and possible transmission of ARG between bacteria.

Thus, to achieve the ultimate turnaround time in a fully automated way and tackle the problem of acquired resistances, we introduce a new method, called TBwDM (Taxonomic Bining with Determinant Mapping) relying on both taxonomic-read binning against a pathogen and ARG Reference DataBase (RDB) followed by assembly. Based on simulated metagenomes, we estimated the performance of our method in terms of ARG recovery, and compared results obtained with MetaCherchant. Finally, we evaluated our method on real bronchoalveolar lavage (BAL) samples.

## Results

### Pipeline TBwDM illustration

Figure 1 illustrates the ability of pipeline TBwDM to retrieve an ARG element that is present (left panel) or absent (right panel) from the pneumonia RDB.

**Figure 1:**
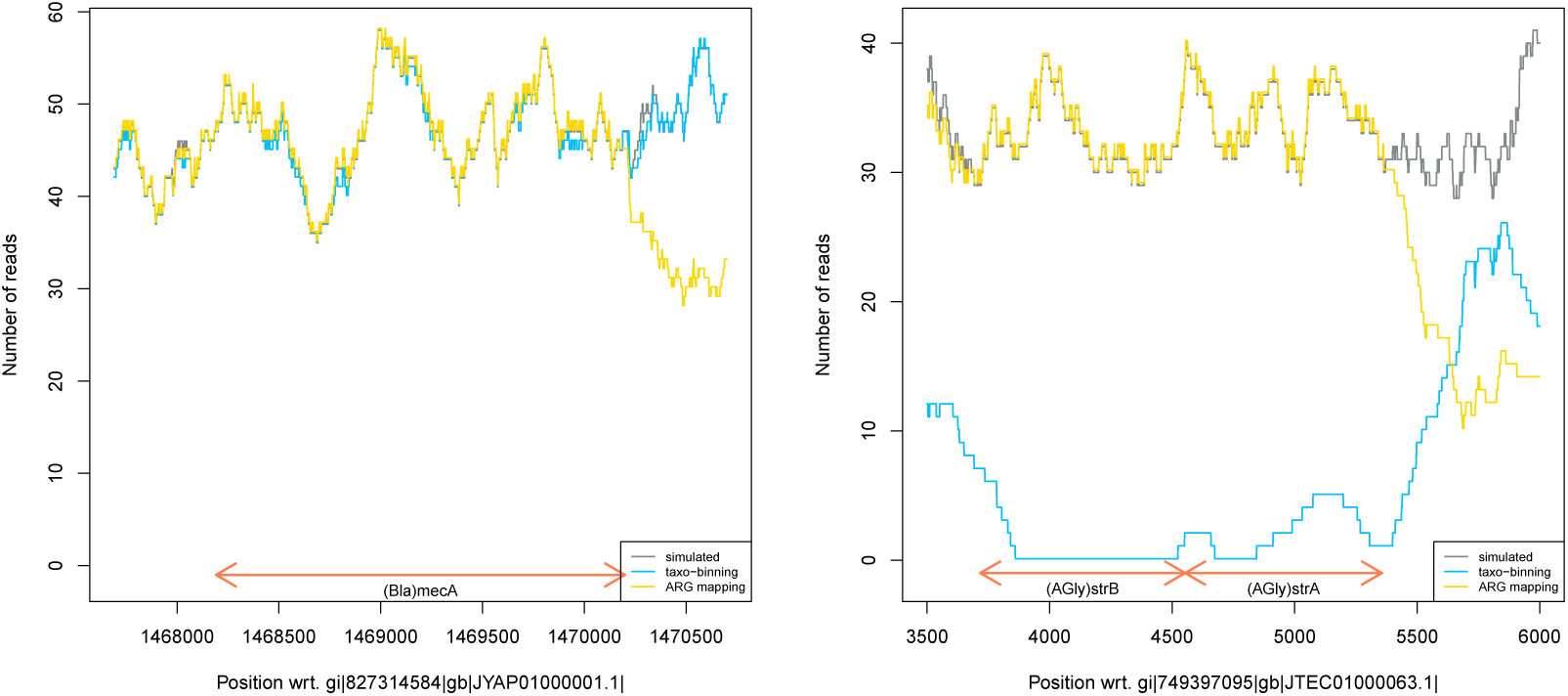
Illustration of pipeline TBwDM in terms of marker recovery. The left panel is centered around the *mecA* gene in *S. aureus* GCF 000013435.1 and the right panel around *strB* and *strA* ARG in *A. baumannii* JTEC01. The grey line corresponds to the number of simulated reads, the blue line to the number of reads retrieved by taxonomic binning against the pneumonia RDB, and the yellow line to the number of reads retrieved by mapping against the ARG RDB.

The left panel is centered around the *mecA* gene in *Staphylococcus aureus* GCF 000013435.1 assembly. *mecA* is a well-described resistance gene that confers resistance to *β*-lactam antibiotics of *S. aureus* strains [19]. *mecA* is present in the representative genomes of *S. aureus* in the pneumonia RDB and also in the ARG RDB, and thus, all the reads simulated in this region were retrieved by both taxonomic binning against the pneumonia RDB and mapping of reads against the ARG RDB (see Figure 1 left panel).

The right panel is centered around the *strA* and *strB* genes in *Acinetobacter baumannii* JTEC01 assembly. These two genes are known to confer resistance to aminoglycosides, and are located in a well-described resistance island (*AbaR*), including several other resistance genes acquired by horizontal gene transfer [20]. These genes are present in genomes from several species in the pneumonia RDB and therefore reads could not be specifically assigned to *A. baumannii*. Thus, as shown in Figure 1, in-silico simulated reads from these two genes were not retrieved by taxonomic binning against the pneumonia RDB (see blue line close to 0 in Figure 1), while they were by mapping against the ARG RDB (see yellow line superimposed with grey line in Figure 1). As a consequence, mapping reads against both ARG RDB and pneumonia RDB allowed us to retrieve all the in-silico simulated reads and to obtain a complete and accurate assembly from the genomic resistance island. Of note, the number of reads obtained by mapping against the ARG RDB decreased slowly outside the resistance gene sequences because paired-end reads were individually mapped with clipping.

### In-silico simulations

#### Species detection

Species detection performance was assessed in terms of precision and recall on confirmed pathogens. As shown in Table 1, both pipelines were extremely precise, with a perfect detection of the 21 pathogens included in the 21 monomicrobial infection simulations, and one false negative among the 43 pathogens included in the 21 polymicrobial infection simulations. The latter error concerned *Enterobacter cloacae complex* JRFQ01, the least abundant of the 3 pathogens included in scenario 11 (see Figure 6), and simulated at a 0.45X coverage. Since this pathogen was correctly detected in monomicrobial infection simulation (coverage 45X) and in scenario 5 of polymicrobial infection simulations (coverages 4.5X and 450X), the detection error in scenario 11 was indicative of the limit of detection of our method, based on post-assembly marker-based taxon confirmation. Detailed confusion matrices are presented in supplementary materials (Supplementary Figures 3 and 4).

**Table 1.** Pathogen detection performance for monomicrobial and polymicrobial infection simulations. The total number of pathogens (# path), the precision/recall for each pipeline, and the total number of false negatives (# FN), if applicable, are provided.

| Pipeline | Monomicrobial |  |  | Polymicrobial |  |  |  |
| --- | --- | --- | --- | --- | --- | --- | --- |
|  | # path. | Prec. | Recall | # path. | # FN | Prec. | Rec. |
| TBo | 21 | 100 | 100 | 43 | 1 | 100 | 97.7 |
| TBwDM | 21 | 100 | 100 | 43 | 1 | 100 | 97.7 |

**Figure 2:**
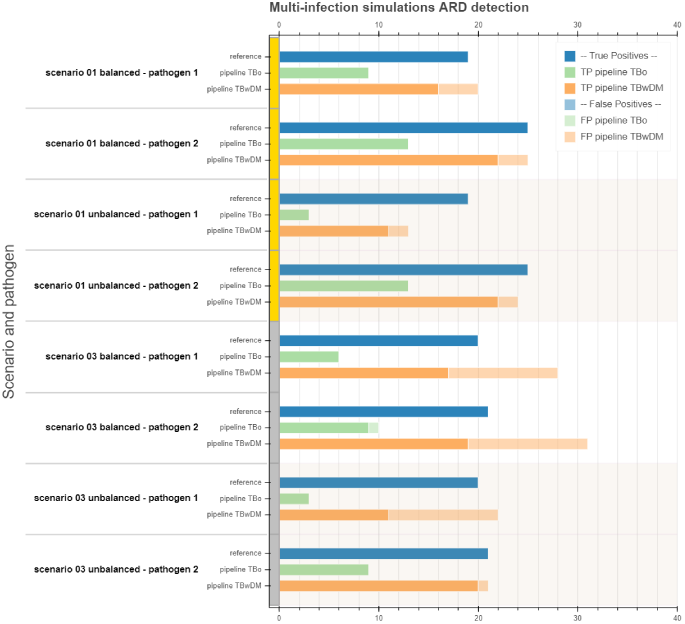
Illustration of polymicrobial simulations ARG detection performance on scenarios 1 and 3, with both balanced and unbalanced coverage configurations. Reference lines state for the true number of ARG in each pathogen. Pipeline TBo and pipeline TBwDM lines show ARG detection results, with solid color standing for true positives, and faded color standing for false positives. False negatives can be deduced by subtracting true positives from the reference. Balanced scenarios simulate 250X coverage for all pathogens, while unbalanced scenarios simulate 4.5X coverage for pathogen 1 and 450X coverage for pathogen 2.

**Figure 3:**
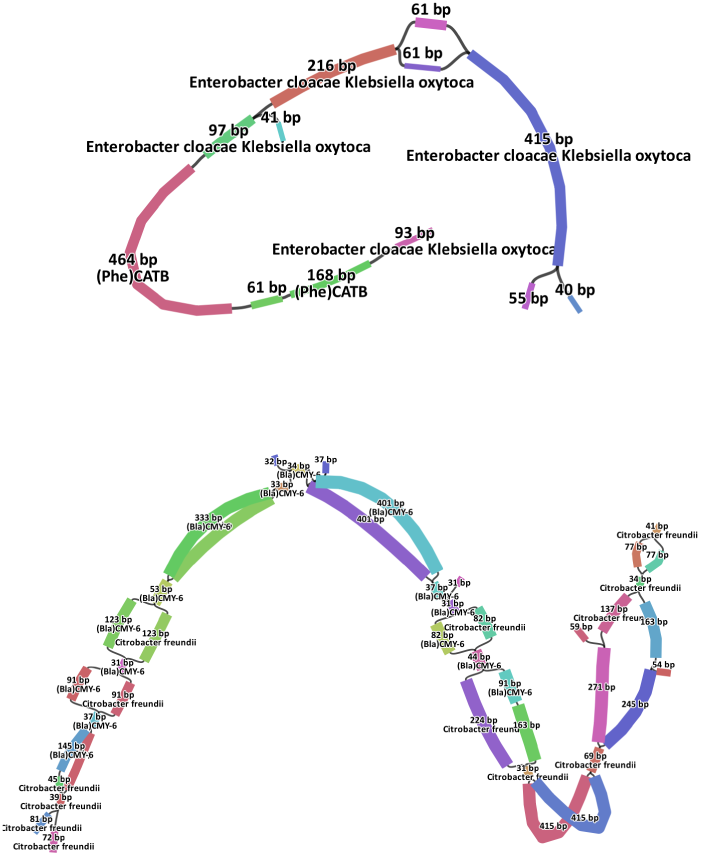
Polymicrobial infection simulations of the ARG genomic context understanding using MetaCherchant. Subgraphs are drawn in the neighborhood of ARG that are detected by our pipeline TBwDM, and related unitigs are blasted against a subset of our pneumonia RDB based on the detected pathogens. Top: scenario 1 unbalanced, ARG *(Phe)CATB* is correctly assigned to both pathogens *E. cloacae complex* and *K. oxytoca*. Bottom: scenario 3 unbalanced, ARG *(Bla)CMY-6* is correctly assigned to *C. freundii*.

**Figure 4:**
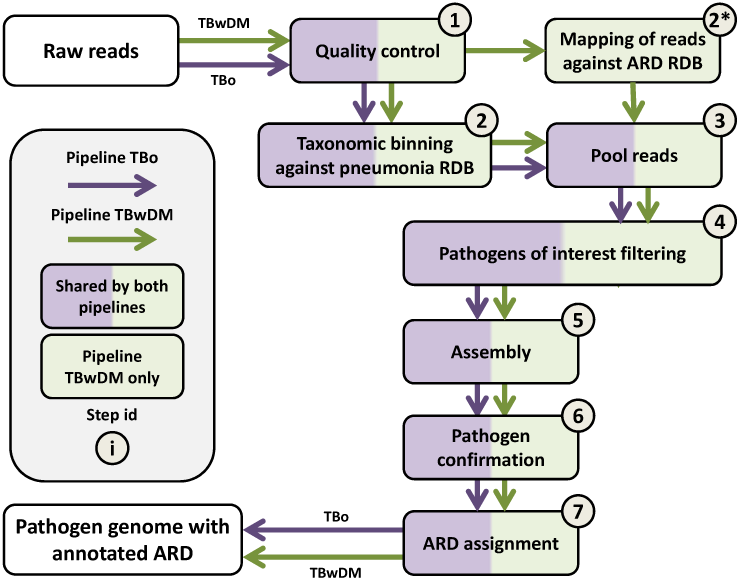
Bioinformatics pipeline for identification of pathogens and ARG from metagenomic sample. In green boxes, new steps are added to pipeline TBwDM to improve ARG recovery. The pneumonia RDB includes genomes from organisms found in the lung and oral cavity.

**Figure 5:**
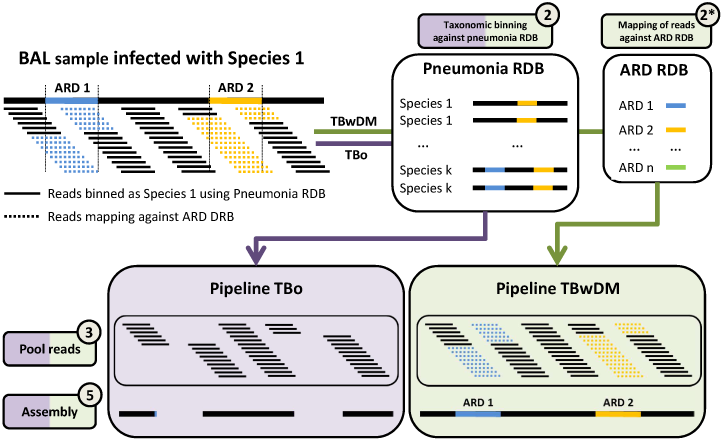
Comparison between pipeline TBo and pipeline TBwDM in terms of ARG recovery.

**Figure 6:**
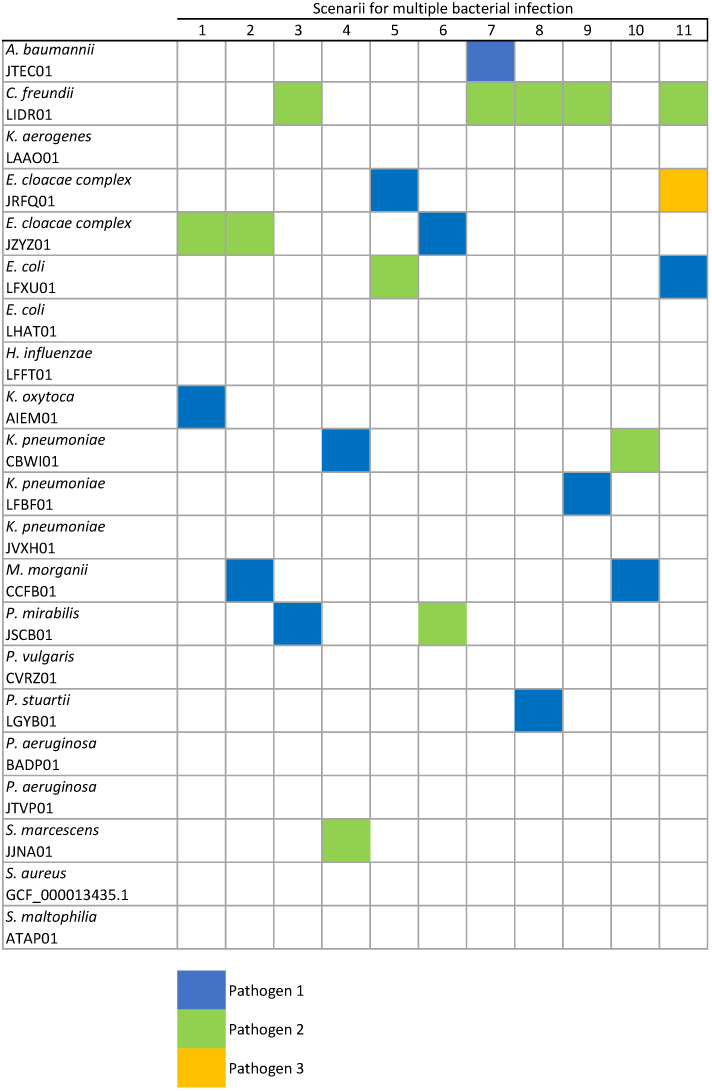
BAL sample simulations. Genomes used for the simulated BAL samples are identified by their NCBI WGS Project identifier, except for *S. aureus* for which we used the NCBI Assembly identifier. Each column corresponds to a scenario of multiple bacterial infection; genomes used for these polymicrobial infection simulations are indicated with a colored square: blue = pathogen 1, green = pathogen 2, and yellow = pathogen 3.

#### Marker recovery

ARG detection and assignment to its correct microbial host was then evaluated for the detected pathogens. For the polymicrobial infection simulations, we used the strategy described in section Bioinformatics pipeline to decide to which pathogen it had to be assigned.

For the 21 monomicrobial infection simulations, as shown in Table 2, both pipelines exhibited high precision (> 97%), while pipeline TBwDM additional step of reads mapping against ARG RDB brought a clear additional value by increasing the recall versus pipeline TBo (Taxonomic Binning only) from 51.5% to 97.3%.

**Table 2.** ARG detection performance for monomicrobial and polymicrobial infection simulations, once ARG are assigned to some pathogens. The total number of pathogens (# path), the total number of ARG (# ARG), and the associated precision/recall are provided.

| Pipeline | Monomicrobial |  |  |  | Polymicrobial |  |  |  |
| --- | --- | --- | --- | --- | --- | --- | --- | --- |
|  | # path. | # ARG | Prec. | Rec. | # path. | # ARG | Prec. | Rec. |
| TBo | 21 | 338 | 98.9 | 51.5 | 42 | 799 | 95.1 | 41.7 |
| TBwDM | 21 | 338 | 97.3 | 97.3 | 42 | 799 | 74.1 | 81.5 |

Regarding the 21 polymicrobial infection simulations, it appeared that there was a significant drop in the performance of both pipelines for ARG detection recall (respectively −10% and −16% vs. monomicrobial infection simulations recall), pipeline TBwDM being impacted in terms of precision as well (− 23% vs. monomicrobial infection simulations precision), see Table 2. However, pipeline TBwDM still achieved better global performance than pipeline TBo, with a larger F1 statistic (77.6% vs. 58.0%). Table 3 illustrates how strongly detection performance depends on pathogen coverage, by focusing on the 2-pathogen’ polymicrobial infection simulations (scenarios 1 to 10, see Figure 6). Indeed, while balanced scenarios (250X coverage for pathogens 1 and 2) brought similar ARG detection performance for both pathogens, unbalanced scenarios led to a drop of recall for the lower-covered pathogen (see pipeline TBwDM, from 89.9% to 54.8%) and an increase of precision for the higher-covered pathogen (see pipeline TBwDM, from 75.5% to 94.8%). The latter can be explained by the fact that unbalanced pathogen coverage disambiguates our ARG assignment step, which uses coverage information to link each ARG to its host.

**Table 3.** ARG detection performance for 2-pathogens polymicrobial infection simulations (scenarios 1 to 10), once ARG are assigned to some pathogens. Path. id refers to pathogen identifier, and # ARG to the total reference number of ARG belonging to pathogens 1 or pathogens 2. Balanced scenarios simulate 250X coverage for all pathogens, while unbalanced scenarios simulate 4.5X coverage for pathogen 1 and 450X coverage for pathogen 2.

| 2-pathogens polymicrobial |  |  | Pipeline TBo |  | Pipeline TBwDM |  |
| --- | --- | --- | --- | --- | --- | --- |
| Scenario | Path. id | # ARG | Precision | Recall | Precision | Recall |
| Balanced | 1 | 168 | 92.4 | 43.5 | 67.4 | 89.9 |
| Balanced | 2 | 208 | 97 | 46.6 | 75.5 | 87.5 |
| Unbalanced | 1 | 168 | 87.8 | 25.6 | 54.8 | 54.8 |
| Unbalanced | 2 | 208 | 99.0 | 47.1 | 94.8 | 87.0 |

Figure 2 illustrates the main outcomes of our experiments, by showing the ARG detection and assignment results obtained on polymicrobial scenarios 1 and 3, both for balanced and unbalanced coverage configurations. It shows that ARG detection recall was improved by pipeline TBwDM, and that unbalanced coverage in 2-pathogens polymicrobial samples improved precision on the higher-covered pathogen (pathogen 2), while it decreased the lower-covered pathogen (pathogen 1) recall at the same time.

We also analyzed ARG detection at the sample level, without the pathogen assignment step (see Table 4). In this settings, the total number of considered ARG decreases from 799 to 693, since 106 ARG are shared by several pathogens included in the same simulated sample. While pipeline TBo performance remained quite stable, pipeline TBwDM precision and recall respectively increased from 74.1% to 94.9% and from 81.5% to 88.5%. These results illustrates the difficulty to correctly assign an ARG to its microbial host, underlying that there is still room for improvement.

**Table 4.** ARG detection performance for polymicrobial simulations, when ARG are considered at the sample level, i.e. <u>not assigned</u> to some pathogens.

| Polymicrobial simulations - at the sample level |  |  |  |  |
| --- | --- | --- | --- | --- |
| Pipeline | # simulations | # ARG | Precision | Recall |
| TBo | 21 | 693 | 97.5 | 44.2 |
| TBwDM | 21 | 693 | 94.9 | 88.5 |
| MetaCherchant | 21 | 693 | NA | 86.7 |

#### Additional insight on detected ARG

In addition to offering pathogen and ARG detection, our bioinformatics pipeline TBwDM can be complemented for understanding genomic context in the neighborhood of detected ARG. To illustrate this, we ran MetaCherchant [18], a graph-based algorithm for extracting ARG and their genomic context (sequence environment) from metagenomic data, on the simulated polymicrobial infections.

Table 4 shows MetaCherchant performance at the sample level (i.e. without pathogen assignment). We can see that MetaCherchant ARG detection recall is roughly the same as for our pipeline TBwDM recall. We did not mention the precision performance on purpose (see NA in Table 4), since pre-processing (e.g. ARG RDB clustering) or post-processing (e.g. best hit selection in each group of redundant subgraphs) should have been designed to reduce false positives and provide a fair estimation.

Then, to come up with both ARG and pathogen identification, so that we can assign ARG to bacterial species, post-processing was carried out to label the neighborhood of the detected ARG in the graph, by blasting the related unitigs (defined as long non-branching paths in the graph) against our pneumonia RDB. This is something that we implemented but, given the length of the aforementionned unitigs (typically a few hundred base pairs) and given the inter-species similarity of our pneumonia RDB (e.g. within Enterobacteriaceae family), perfect matches could occur between unitigs and a set of different species, which made unitigs assignation and pathogen identification impossible.

### Real data

Sample 1 contained *Escherichia coli* species, while samples 2 et 3 were polymicrobial samples with *Klebsiella pneumoniae* and *Haemophilus influenzae* (sample 2), and *E. coli* and *Klebsiella aerogenes* (sample 3). The two pipelines detected the same pathogens, and results were in accordance with standard microbiological culture. Performance for ARG detection for the two pipelines are presented in Table 5. Briefly, the average recall over the 5 isolates was 95.8% with TBwDM and 60.4% with TBo, the average precision was 83.3% with TBwDM and 76.3% with TBo. These results were coherent with the ones obtained in simulation studies, i.e. a better performance with TBwDM than with TBo. Details about the ARG marker detected in the strain sequences, and by the two pipelines can be found in supplementary Figure 11. As can be seen in this figure, for sample_2, both TBo and TBwDM failed to uniquely assign *(Elf)TUFAB* elongation factor to *K. pneumoniae*; for sample_3, TBwDM failed to uniquely assign *(Sul)Sul2* and *(Tet)TetR* to *E. coli*, leading in both cases to a decreased precision. This is because we conservatively chose to assign ARG markers to all the detected pathogens in case of ambiguous assignments (see Methods).

**Table 5.**
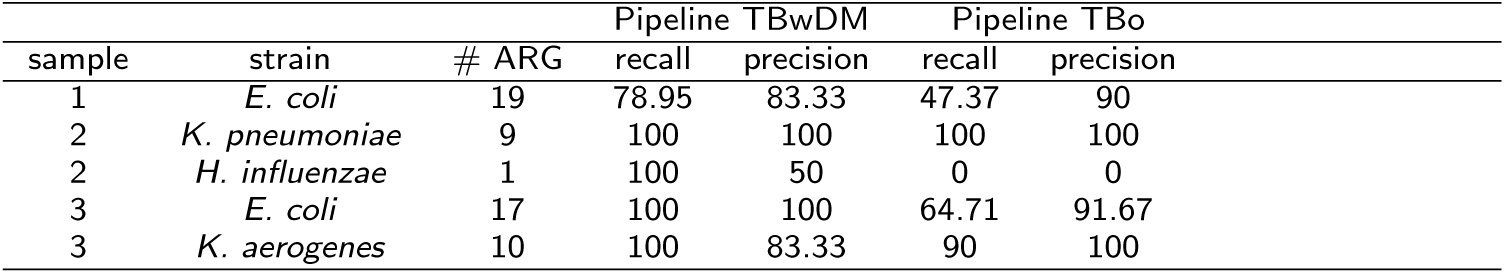
ARG detection performance on real BAL samples (# ARG refers to the number of ARG found in the strain sequence).

| sample | strain | # ARG | Pipeline TBwDM |  | Pipeline TBo |  |
| --- | --- | --- | --- | --- | --- | --- |
|  |  |  | recall | precision | recall | precision |
| 1 | *E. coli* | 19 | 78.95 | 83.33 | 47.37 | 90 |
| 2 | *K. pneumoniae* | 9 | 100 | 100 | 100 | 100 |
| 2 | *H. influenzae* | 1 | 100 | 50 | 0 | 0 |
| 3 | *E. coli* | 17 | 100 | 100 | 64.71 | 91.67 |
| 3 | *K. aerogenes* | 10 | 100 | 83.33 | 90 | 100 |

## Discussion/Conclusion

In this article, we presented a new bioinformatics pipeline to detect pathogens and their ARG in BAL from patients with HAP-VAP. To improve pathogen genome assembly and ARG recall, we performed read mapping against an ARG RDB in parallel to taxonomic read binning. We used a machine learning algorithm [21] to classify reads into human, flora (commensal oral bacteria), or pneumonia-related pathogens with a reference database optimized for BAL sample metagenomes. Moreover, we optimized read mapping parameters to allow clipping and thus mapping reads at the junction between the ARG and its chromosomal environment with a good sensitivity. In the case of in-silico simulated monomicrobial infections, these two optimizations allowed us to reach very good performance for pathogen detection (100% precision and recall) and ARG detection (97.3% precision and 97.3% recall). In the case of simulated polymicrobial infections, we leveraged parallel taxonomic binning and ARG read mapping to define assignment rules and thus link ARG to host pathogens. This step is valuable in the context of clinical metagenomics [4] because it allows the microbiologist and the clinician to prescribe a better therapy. The precision of pipeline TBwDM (resp. pipeline TBo) was 74.1% (resp. 95.1%) and the recall 81.5% (resp. 41.7%), leading to a larger F1 statistic for pipeline TBwDM (77.6%) compared to pipeline TBo (58.0%). As expected, performance results were dependent on the sequencing depth of the pathogen, with much better results for highly covered pathogens. Moreover, in real BAL samples (monomicrobial and polymicrobial samples), both recall and precision for ARG detection were higher with the combined approach than with the standard one (35 points difference for recall, and 7 points difference for precision)

Here, we defined empiric rules, based on contextual detection and coverage level, to link an ARG to its host pathogen when the ARG is located on the chromosome. However, assigning an ARG located on a plasmid to the correct host pathogen still remains a challenge. Using coverage information, and relying on the hypothesis that an ARG located on a plasmid should have the same or a greater coverage level than the main chromosome would require that plasmids and chromosomes are extracted and sequenced with the same efficiency (see section Bioinformatics pipeline), and this hypothesis was not confirmed by [22]. Thus, in absence of confirmation that the ARG was located on a chromosome, the ARG was conservatively assigned to all the pathogens present in the sample.

We also positioned MetaCherchant software [18] as a valuable post-processing tool for our bioinformatics pipeline, allowing to obtain a reconstruction of the genomic context around detected ARG. While our original intention was to benchmark MetaCherchant with our pipelines, it could not directly compete in terms of ARG detection precision given the high similarity between ARG sequences in our ARG RDB. Indeed, when it was directly executed on the simulated polymicrobial infections reads, it provided a lot of false positives by drawing redundant ARG sub-graphs. This is a well-known limitation of resistome profiling tools in metagenomics, one solution being to remove ARG redundancy by clustering or indexing techniques. However, it proved useful when restricted to the set of pathogens and ARG detected by our pipeline. Indeed, as suggested in [23], it was able to provide some valuable insight on the genomic environment of the ARG which were detected by TBwDM. What we suggest is to focus on these detected ARG, and to label their subgraphs by blasting the related unitigs against a subset of our pneumonia RDB, by restricting it to the detected pathogens. As an example, Figure 3 illustrates two different genomic contexts that we met in our simulations: one related to some ARG that was correctly assigned to one pathogen, and one related to some ARG that was correctly assigned to both pathogens in the simulated sample. Additional graphs are available in the supplementary materials, including examples of cases of false assignment.

Currently, pipeline TBwDM provides a list of ARG with assignment to their host pathogens and requires the guidance of a highly trained molecular microbiologist to interpret the link between presence/absence of ARG and antibiotic resistance. A major improvement of this pipeline would be the automated prediction of antibiotic susceptibility/resistance profile for each detected pathogen. Although such prediction from isolates is a very active field of research with clinically useful performance achieved for *Mycobacterium tuberculosis* [24], *S. aureus* [25] [26], *K. Pneumoniae* [27] or *E. coli* [28] among others, translating it to a metagenomic samples remains a challenge.

Another great challenge would be the absolute quantification of pathogens with the reporting of genome copy numbers into interpretable colony forming unit figures actionable by the microbiologist. Indeed, distinguishing between colonization and infection when a pathogen is detected, is necessary to initiate a possible drug treatment [29].

Finally, a current limitation of our process for its introduction into clinical routine is the sequencing turnaround time. Indeed, the sequencing time with the Illumina^®^MiSeq sequencer is currently around 39 hours (2 × 250 nucleotides paired-end) which greatly limits its usefulness in routine settings. An alternative strategy would be to combine our pipeline with real-time and long-read sequence data as those provided by the Oxford Nanopore Minion, as recently suggested in [30], where authors consider that it would be necessary to associate ARG to organisms in Nanopore metagenomics applied to bacterial lower respiratory infection.

## Abbreviations

HAP-VAP: Hospital-Acquired and Ventilator-Associated Pneumonia
DNA: Desoxyribose Nucleic Acid
BAL: BronchoAlveolar Lavage
ARG: Antibiotic Resistance Gene
RDB: Reference DataBase
TBwDM: Taxonomic Binning with Determinant Mapping
TBo: Taxonomic Binning only

## Methods

In this section we describe the bioinformatic pipelines that we developed to identify HAP-VAP pathogens with their resistance determinants from BAL samples. To deal with our pneumonia application, we decided to focus on the 20 most relevant pathogens for HAP-VAP infections [31], 18 Gram-negative and 2 Gram-positive bacterial species, which we list hereafter: *Escherichia coli, Enterobacter cloacae, Klebsiella aerogenes*^[1]^, *Klebsiella oxytoca, Klebsiella pneumoniae, Citrobacter koseri, Citrobacter freundii, Morganella morganii, Proteus mirabilis, Proteus vulgaris, Providencia stuartii, Serratia marcescens, Hafnia alvei, Acinetobacter baumannii, Haemophilus influenzae, Legionella pneumophila, Pseudomonas aeruginosa, Stenotrophomonas maltophilia, Staphylococcus aureus* and *Streptococcus pneumoniae*.

We also describe the datasets used, both simulated and read datasets as well as performance evaluation criteria.

### Bioinformatics pipeline

The classical metagenomic approach to identify pathogens and ARG in a sample requires the following steps: reads quality control, trimming and filtering of poor quality reads, elimination of host (human) DNA, taxonomic read binning using a reference database (RDB) including genomes from the organisms found in the region of interest (in our application, a “pneumonia” RDB which focuses on lung and oral cavity), assembly of reads corresponding to each pathogen present in the sample into contigs, and finally annotation of contigs with respect to an ARG RDB [32, 33]. This bioinformatics pipeline, called pipeline TBo (Taxonomic Binning only), is illustrated in Figure 4 (see steps 1 to 7, following purple arrows). The output of this pipeline is the set of pathogen(s) present in the sample with the set of ARG contained in their genomes.

As can be seen in Figure 5 (left-side), the main drawback of pipeline TBo is its lack of sensitivity for ARG detection. As an example, let’s assume: 1/ a BAL sample containing a strain from Species 1 and harboring 2 ARG (see blue and orange segments in Figure 5); 2/ a pneumonia RDB containing representative sequences from Species 1, none of these sequences harboring ARG 1. In such a situation, as can be seen in Figure 5, pipeline TBo only retrieves the reads outside ARG 1 for assembly and ARG 1 is missing from the final assembly. Another situation where pipeline TBo lacks sensitivity is illustrated with ARG 2 (see orange segment in Figure 5). ARG 2 is present in the genomes from 2 different species (species 1 and *k* in Figure 5) and thus cannot be specifically assigned to a single species. In this case, pipeline TBo only retrieves the reads outside ARG 2 for assembly, and ARG 2 is missing from the final assembly. This is typically the case of an ARG which has recently been acquired through horizontal gene transfer by a pathogen facing selective pressure [34, 35]: it can be either absent from the reference genomes of this pathogen (case of a recent acquisition) or present in reference genomes of several species. This problem is expected to be frequent in our context because several of the bacteria involved in the HAP-VAP infections are known to resort to horizontal gene transfer to rapidly improve their fitness, e.g. *P. aeruginosa* [36], *K. pneumoniae* [37], *A. baumannii* [38], or *S. aureus* [39].

To improve ARG assignment to their host pathogens, we developed a new pipeline, called TBwDM (Taxonomic Binning with Determinant Mapping), with an additional step of read mapping against the ARG RDB (see step 2* in Figure 4) performed in parallel of the taxonomic binning step (step 2). Then, reads for each pathogen are pooled with the reads mapped against the ARG RDB (step 3), and all reads are assembled together (step 5) after filtering (step 4). As illustrated in Figure 5, even if ARG 1 is absent from the representative sequences of Species 1, reads falling in this region are retrieved by mapping against the ARG RDB. Moreover, even if ARG 2 is harbored by several species and thus ARG reads cannot be classified at the species level, reads falling in this region can be retrieved by mapping against the ARG RDB. All reads are then pooled together and the final assembly includes ARG 1 and ARG 2.

For the taxonomic read binning step, we relied on a multi-class predictor [21] that allowed us to classify each read into 22 classes: the 20 classes for the afore-mentioned 20 pathogens of interest, a human class, and a flora class. Indeed, BAL samples typically include human cells but also commensal bacteria from the oral cavity [40]. Bacteria included in the flora class were selected based on literature [41, 42], results from the Human Microbiome Project [43], but also on internally sequenced pathogen-free BAL samples. A Krona chart of the bacterial composition of the flora class is available in Supplementary Figure 1. Reads classified as human or flora were filtered to focus only on the reads which have been assigned to the pathogens of interest. To train the multi-class predictor, we built a pneumonia RDB including bacterial genomes from the 20 pathogens of interest, human genome (version GRCh37), and genomes from the flora class. These genomes are both public (e.g. Patric, RefSeq, FDA ARGOS) and private (strains sequences from BAL samples). The median number of pathogen genomes per species was 42 with inter-quartile-range equal to [19; 297], and the total number of genomes in the flora class was 7593. The multi-class model was a linear penalized predictor based on the genome frequency of 12-mers [21].The performance of the predictor on simulated strains absent from the pathogens RDB is reported in Supplementary Figure 2. It has to be noticed that we experimented taxonomic binning with other tools, such as Kraken [7], and that we obtained similar results.

For mapping reads against the ARG RDB, we relied on the bwa mem program (bwa 0.7.8 version) [44]. Mapping parameters were optimized as described in [45] to maximize the sensitivity of reads retrieval, in particular for reads at the junction between the ARG and the genome: `bwa mem -a -T 0 -k 16 -L 5 -d 100`. Moreover, we used single-end reads mapping, even for paired-end reads and pooled both reads of the pair for assembly, as soon as one read of the pair mapped. The ARG RDB was built from the DBGWAS ARG RDB [46]. We removed from this database perfectly identical sequences and also markers corresponding to the main efflux families (ABC, MFS, …), as well as intrinsic genes (topoisomerase IV subunits, gyrase) so that only ARG whose presence/absence is indicative of antibiotic resistance are used.

Then, reads binned as pathogen and reads mapped against the ARG RDB were pooled together for assembly. In case of polymicrobial infection, reads mapped against the ARG RDB were pooled with reads of each pathogen. All the pathogens with average coverage > 1 were then assembled. Assembly was performed using `idba_ud500` assembler (idba ud 1.1.1) [47], with following parameters: `--mink 40 --maxk 250 --min_pairs 2`.

In the pathogen confirmation step (see step 6 in Figure 4), we used the blast algorithm to align the assembly against a pathogen marker database, built by selecting MetaPhlAn2 markers [11] corresponding to the 20 pathogens of interest. These markers were selected to be clade-specific and allow unambiguous taxonomic assignments. This step was added to avoid spurious pathogen detection due to erroneous binning. The blast algorithm was run with coverage = 75% and percent identity = 97%. Whenever a marker of the tested pathogen was detected, the pathogen was confirmed. If a marker from another pathogen was detected, the pathogen was not confirmed. When no pathogen marker was detected (typically observed for partial assembly due to low coverage), the pathogen was flagged as dubious.

Finally, we used blast to align the assembly of any confirmed pathogen against the ARG RDB, with coverage and percent identity parameters equal to 80%. In case of polymicrobial infection, it was particularly important to correctly assign each ARG to each pathogen (see step 7 in Figure 4). We used the following rules to link each ARG to its host pathogen: we checked whether > 5% of the reads mapped against the contig containing the ARG have been retrieved by taxonomic binning against the pneumonia RDB (see Figure 4), and in parallel, we checked that the median coverage of the marker was > 1/3 to the median coverage of the pathogen. If yes, this means that the ARG was integrated in the genome of the pathogen and should be assigned to it, otherwise the marker was assigned to all the detected pathogens for further investigation. Thresholds were set empirically, being conservative enough to avoid missing ARG.

### Data

In the following sections, we describe both simulated and real datasets along with the indicators used to evaluate the performance of our pipelines.

#### Monomicrobial infection simulations: BAL samples containing a single pathogen

The purpose of this dataset was to evaluate the ability of the pipeline to identify the correct bacterial species and retrieve the whole set of ARG contained in their genome. Thus, we selected 21 strains with publicly available genomes (see Figure 6 for NCBI WGS project identifier) from the pathogens of interest in HAP-VAP infections, based on their ARG contents. We simulated reads from each of these genomes, as well as flora reads from bacterial genomes of the flora class (see Methods above) to create in silico simulated BAL samples containing a single pathogen species. We simulated 300,000 reads of pathogens (equivalent to a 45X average genome coverage) and 150,000 reads of flora. We simulated 2 × 300 nucleotides paired-end reads, assuming a Illumina MiSeq error profile.

#### Polymicrobial infection simulations: BAL samples containing several pathogens

The purpose of this dataset was to evaluate the ability of the pipeline to retrieve the whole set of ARG and to correctly assign each ARG to each pathogen. Figure 6 presents the 11 evaluated scenarios; for each scenario, pathogen 1 is indicated by a blue box, pathogen 2 by a green box and, if applicable, pathogen 3 is marked by a yellow box. For each scenario (except scenario 11), we created 2 simulated metagenomes, a “balanced” dataset with an equal quantity of pathogen 1 and 2 (average coverage equal to 250X), and a ‘unbalanced” dataset with a pathogen 1 average coverage equal to 4.5X and pathogen 2 equal to 450X. For scenario 11, the average coverage of pathogen 1 was 45X, the average coverage of pathogen 2 was 450X, and the average coverage of pathogen 3 was 0.45X. As for simulations with a single pathogen, flora reads were also included, with an average coverage equal to 0.45X.

In total, we evaluated our bioinformatics pipelines on a set of 21 polymicrobial infection simulations (scenarios 1 to 10 with 2 coverage configurations, and scenario 11), involving 43 pathogens (20 simulations with 2 pathogens, and 1 with 3 pathogens).

#### Real BAL samples

We also assessed the performance of the pipelines on real samples, using 3 BAL samples from patients in intensive-care units. We sequenced the BAL samples (sample-level characterization) and the strains isolated from the BAL samples (strain-level characterization), and compared the ARG detected by the pipelines on the BAL samples to the ARG detected on the isolates (gold-standard). For the sample-level characterization, 600 *µ*L of BAL sample was treated as described in [48] to favor the differential lysis of human cells and thus reduce the quantity of human DNA before microbial DNA extraction. Nucleic acids were extracted using the bioMérieux easyMAG^®^instrument. Libraries were prepared using the Nextera^®^XT DNA Library Preparation Kit, and sequenced with the Illumina^®^MiSeq instrument (2 × 250 nucleotides paired-end sequencing with V3 chemistry). For the strain-level characterization, strains were isolated from the BAL samples by culture. Nucleic acids were extracted using the Qiagen^®^DNeasY UltraClean Microbial Kit, and sequenced with the Illumina^®^MiSeq instrument (2 × 200 nucleotides paired-end sequencing with V3 chemistry).

### Evaluation criteria

Because our goal was to estimate how well pipelines were able to detect pathogens in metagenomic samples, and how many ARG could be retrieved and assigned to the right pathogen, our evaluation strategy was twofold.

The performance of pathogen detection was first assessed in terms of precision and recall. In this context, precision referred to the fraction of detected pathogens that were actually present in the related sample, and recall referred to the fraction of sampled pathogens that were detected by our pipelines. While getting the reference pathogen was straightforward for the simulated data, we used traditional microbiological culture to get the reference pathogen in the real BAL samples. Only confirmed pathogens (see section Bioinformatics pipeline) were considered. For the specific cases of the polymicrobial infection simulations, we evaluated how much the detection recall was affected by pathogen coverage.

Then, in order to distinguish the pathogen detection performance from the ARG detection and assignment performance, the latter was only assessed for the correctly detected pathogens. This means that ARG belonging to undetected pathogens were not included in the performance calculation. For simulated data, reference ARG were obtained by blasting the 21 public genomes against our ARG RDB, while for real data, reference ARG were obtained by blasting the sequences of the strains isolated from the BAL samples against our ARG RDB.

We also illustrated how we can get some additional insight on the detected ARG genomic environments by using MetaCherchant [18], which is a graph-based algorithm for extracting ARG and their genomic context from metagenomic data. We evaluated how MetaCherchant deals with ARG detection from the raw simulated polymicrobial infection reads, and how we could favorably use this tool as a post-processing of our bioinformatics pipeline. As proposed in https://github.com/ctlab/metacherchant/, we used the single-metagenome mode with the default parameterization: `metacherchant.sh --tool environment-finder --k 31 --coverage=5 --maxkmers=100000`. All the graphs representing ARG and their genomic environments were generated using Bandage visualization tool (available at https://rrwick.github.io/Bandage/).

## Declarations

### Ethics approval and consent to participate

Three LBA samples from patients hospitalized in Intensive Care Unit were collected by the hospital of Annecy (France) for diagnostic purpose. Well characterized left-over LBA samples were then proposed for research activities. As mentioned in the 3° of the L.1121-1 article of the French Public Health Code, this study was classified as a non-interventional research. This study was registered on the database of the National Drug Safety Agency (ANSM) with the identification number 2017-A00253. Governed by the 1978 data protection law and particularly by the reference methodology n°3, deliberation n°2016-263 of July 21th, 2016, an oral and written individual information was delivered by the clinician to each patient. Oral non-opposition obtained from each patient was documented in a specific form and signed by the clinician, archived in the medical file. Conformingly to the French regulation and ethical requirements, the study protocol, an information note and the non-opposition form were approved by the Ethical Committee, Sud Mediterranean I, Sainte Marguerite hospital, 270 Bd Sainte Marguerite, 13274 MARSEILLE, with reference number IORG0009097. The approval was obtained on April 05th, 2017 under the project reference number 17 19.

### Consent for publication

Not applicable

### Author’s contributions

MT, ER, GP, SS, GGu, and PM developped the bioinformatics pipeline and performed the analysis on simulated and real data. VL, SH, CM, GGe, and JS conceived and designed the study, and with AL and KL collected the data.

MT, GP, and SS wrote the article and all the authors reviewed it.

## Acknowledgements

The authors would like to thank Alex van Belkum, Scientific Director of Microbiology Research at bioMérieux, and Bruno Lacroix, head of the Data Science department at bioMérieux, for proofreading this article.

## Availability of data and materials

The data that support the findings of this study are available from the corresponding author upon reasonable request.

## Footnotes

[1] previously known as *Enterobacter aerogenes*.

## References

1. Naccache, S.N., Federman, S., Veeeraraghavan, N., Zaharia, M., Lee, D., Samayoa, E., Bouquet, J., Greninger, A.L., Luk, K.-C., Enge, B., et al.: A cloud-compatible bioinformatics pipeline for ultrarapid pathogen identification from next-generation sequencing of clinical samples. Genome research (2014)

2. Minot, S.S., Krumm, N., Greenfield, N.B.: One codex: a sensitive and accurate data platform for genomic microbial identification. BioRxiv, 027607 (2015)

3. Ruppé, E., Baud, D., Schicklin, S., Guigon, G., Schrenzel, J.: Clinical metagenomics for the management of hospital-and healthcare-acquired pneumonia. Future microbiology 11 (3), 427–439 (2016). doi:10.2217/fmb.15.144

4. Ruppé, E., Cherkaoui, A., Lazarevic, V., Emonet, S., Schrenzel, J.: Establishing Genotype-to-Phenotype Relationships in Bacteria Causing Hospital-Acquired Pneumonia: A Prelude to the Application of Clinical Metagenomics. Antibiotics (2017). doi:10.3390/antibiotics6040030

5. Cassini, A., Högberg, L.D., Plachouras, D., Quattrocchi, A., Hoxha, A., Simonsen, G.S., Colomb-Cotinat, M., Kretzschmar, M.E., Devleesschauwer, B., Cecchini, M., et al.: Attributable deaths and disability-adjusted life-years caused by infections with antibiotic-resistant bacteria in the eu and the european economic area in 2015: a population-level modelling analysis. The Lancet Infectious Diseases 19 (1), 56–66 (2019)

6. Piskin, N., Aydemir, H., Oztoprak, N., Akduman, D., Comert, F., Kokturk, F., Celebi, G.: Inadequate treatment of ventilator-associated and hospital-acquired pneumonia: risk factors and impact on outcomes. BMC infectious diseases 12 (1), 268 (2012)

7. Wood, D.E., Salzberg, S.L.: Kraken: Ultrafast metagenomic sequence classification using exact alignments. Genome Biology (2014). doi:10.1186/gb-2014-15-3-r46

8. Langmead, B., Salzberg, S.L.: Fast gapped-read alignment with Bowtie 2. Nature methods (2012). doi:10.1038/nmeth.1923. #14603

9. Kim, D., Song, L., Breitwieser, F.P., Salzberg, S.L.: Centrifuge: Rapid and sensitive classification of metagenomic sequences. Genome Research (2016). doi:10.1101/gr.210641.116

10. Huson, D.H., Beier, S., Flade, I., Górska, A., El-Hadidi, M., Mitra, S., Ruscheweyh, H.J., Tappu, R.: MEGAN Community Edition - Interactive Exploration and Analysis of Large-Scale Microbiome Sequencing Data. PLoS Computational Biology (2016). doi:10.1371/journal.pcbi.1004957

11. Truong, D.T., Franzosa, E.A., Tickle, T.L., Scholz, M., Weingart, G., Pasolli, E., Tett, A., Huttenhower, C., Segata, N.: MetaPhlAn2 for enhanced metagenomic taxonomic profiling. Nature Methods 12 (10), 902 (2015). doi:10.1038/nmeth.3589

12. Nguyen, N.P., Mirarab, S., Liu, B., Pop, M., Warnow, T.: TIPP: Taxonomic identification and phylogenetic profiling. Bioinformatics (2014). doi:10.1093/bioinformatics/btu721

13. Sczyrba, A., Hofmann, P., Belmann, P., Koslicki, D., Janssen, S., Dröge, J., Gregor, I., Majda, S., Fiedler, J., Dahms, E., Bremges, A., Fritz, A., Garrido-Oter, R., Jørgensen, T.S., Shapiro, N., Blood, P.D., Gurevich, A., Bai, Y., Turaev, D., Demaere, M.Z., Chikhi, R., Nagarajan, N., Quince, C., Meyer, F., Balvočiutė, M., Hansen, L.H., Sørensen, S.J., Chia, B.K.H., Denis, B., Froula, J.L., Wang, Z., Egan, R., Don Kang, D., Cook, J.J., Deltel, C., Beckstette, M., Lemaitre, C., Peterlongo, P., Rizk, G., Lavenier, D., Wu, Y.W., Singer, S.W., Jain, C., Strous, M., Klingenberg, H., Meinicke, P., Barton, M.D., Lingner, T., Lin, H.H., Liao, Y.C., Silva, G.G.Z., Cuevas, D.A., Edwards, R.A., Saha, S., Piro, V.C., Renard, B.Y., Pop, M., Klenk, H.P., Göker, M., Kyrpides, N.C., Woyke, T., Vorholt, J.A., Schulze-Lefert, P., Rubin, E.M., Darling, A.E., Rattei, T., McHardy, A.C.: Critical Assessment of Metagenome Interpretation - A benchmark of metagenomics software. Nature Methods (2017). doi:10.1038/nmeth.4458

14. Li, D., Luo, R., Liu, C.M., Leung, C.M., Ting, H.F., Sadakane, K., Yamashita, H., Lam, T.W.: MEGAHIT v1.0: A fast and scalable metagenome assembler driven by advanced methodologies and community practices (2016). doi:10.1016/j.ymeth.2016.02.020

15. Alneberg, J., Bjarnason, B.S., De Bruijn, I., Schirmer, M., Quick, J., Ijaz, U.Z., Lahti, L., Loman, N.J., Andersson, A.F., Quince, C.: Binning metagenomic contigs by coverage and composition. Nature Methods (2014). doi:10.1038/nmeth.3103

16. Wu, Y.W., Simmons, B.A., Singer, S.W.: MaxBin 2.0: An automated binning algorithm to recover genomes from multiple metagenomic datasets. Bioinformatics (2015). doi:10.1093/bioinformatics/btv638

17. Quince, C., Walker, A.W., Simpson, J.T., Loman, N.J., Segata, N.: Shotgun metagenomics, from sampling to analysis (2017). doi:10.1038/nbt.3935

18. Olekhnovich, E.I., Vasilyev, A.T., Ulyantsev, V.I., Kostryukova, E.S., Tyakht, A.V.: Metacherchant: analyzing genomic context of antibiotic resistance genes in gut microbiota. Bioinformatics 34 (3), 434–444 (2018). doi:10.1093/bioinformatics/btx681

19. Hiramatsu, K., Ito, T., Tsubakishita, S., Sasaki, T., Takeuchi, F., Morimoto, Y., Katayama, Y., Matsuo, M., Kuwahara-Arai, K., Hishinuma, T., et al.: Genomic basis for methicillin resistance in staphylococcus aureus. Infection & chemotherapy 45 (2), 117–136 (2013)

20. Pagano, M., Martins, A.F., Barth, A.L.: Mobile genetic elements related to carbapenem resistance in acinetobacter baumannii. brazilian journal of microbiology 47 (4), 785–792 (2016)

21. Vervier, K., Mahé, P., Tournoud, M., Veyrieras, J.-B., Vert, J.-P.: Large-scale machine learning for metagenomics sequence classification. Bioinformatics 32 (7), 1023–1032 (2015)

22. Ruppé, E., Lazarevic, V., Girard, M., Mouton, W., Ferry, T., Laurent, F., Schrenzel, J.: Clinical metagenomics of bone and joint infections: a proof of concept study. Scientific reports 7 (1), 7718 (2017)

23. Rowe, W.P.M., Winn, M.D.: Indexed variation graphs for efficient and accurate resistome profiling. Bioinformatics 34 (21), 3601–3608 (2018). doi:10.1093/bioinformatics/bty387

24. Coll, F., McNerney, R., Preston, M.D., Guerra-Assunção, J.A., Warry, A., Hill-Cawthorne, G., Mallard, K., Nair, M., Miranda, A., Alves, A., et al.: Rapid determination of anti-tuberculosis drug resistance from whole-genome sequences. Genome medicine 7 (1), 51 (2015)

25. Bradley, P., Gordon, N.C., Walker, T.M., Dunn, L., Heys, S., Huang, B., Earle, S., Pankhurst, L.J., Anson, L., De Cesare, M., et al.: Rapid antibiotic-resistance predictions from genome sequence data for staphylococcus aureus and mycobacterium tuberculosis. Nature communications 6, 10063 (2015)

26. Mahé, P., Tournoud, M.: Predicting bacterial resistance from whole-genome sequences using k-mers and stability selection. BMC bioinformatics 19 (1), 383 (2018)

27. Nguyen, M., Brettin, T., Long, S.W., Musser, J.M., Olsen, R.J., Olson, R., Shukla, M., Stevens, R.L., Xia, F., Yoo, H., et al.: Developing an in silico minimum inhibitory concentration panel test for klebsiella pneumoniae. Scientific reports 8 (1), 421 (2018)

28. Moradigaravand, D., Palm, M., Farewell, A., Mustonen, V., Warringer, J., Parts, L.: Prediction of antibiotic resistance in escherichia coli from large-scale pan-genome data. PLoS computational biology 14 (12), 1006258 (2018)

29. Simner, P.J., Miller, S., Carroll, K.C.: Understanding the Promises and Hurdles of Metagenomic Next-Generation Sequencing as a Diagnostic Tool for Infectious Diseases. Clinical Infectious Diseases 66 (5), 778–788 (2017). doi:10.1093/cid/cix881. http://oup.prod.sis.lan/cid/article-pdf/66/5/778/23879884/cix881.pdf

30. Charalampous, T., Kay, G.L., Richardson, H., et al.: Nanopore metagenomics enables rapid clinical diagnosis of bacterial lower respiratory infection. Nature Biotechnology 37, 783–792 (2019)

31. Barbier, F., Andremont, A., Wolff, M., Bouadma, L.: Hospital-acquired pneumonia and ventilator-associated pneumonia: recent advances in epidemiology and management. Current opinion in pulmonary medicine 19 (3), 216–228 (2013)

32. Guigon, G., Santiago-Allexant, E., Lanet, V., Bonnaud, B., Le Priol, C., Tournoud, M., Gervasi, G., Schrenzel, J., Mallard, F., Veyrieras, J.B.: ECCMID. In: Pathogen Characterization Within the Microbial Flora of Bronchoalveolar Lavages by Direct Sample Sequencing (2015)

33. Breitwieser, F.P., Lu, J., Salzberg, S.L.: A review of methods and databases for metagenomic classification and assembly. Briefings in bioinformatics (2017)

34. Lerminiaux, N.A., Cameron, A.D.: Horizontal transfer of antibiotic resistance genes in clinical environments. Canadian journal of microbiology (999), 1–11 (2018)

35. Ochman, H., Lawrence, J.G., Groisman, E.A.: Lateral gene transfer and the nature of bacterial innovation. nature 405 (6784), 299 (2000)

36. Kung, V.L., Ozer, E.A., Hauser, A.R.: The accessory genome of pseudomonas aeruginosa. Microbiology and molecular biology reviews 74 (4), 621–641 (2010)

37. Wyres, K.L., Holt, K.E.: Klebsiella pneumoniae as a key trafficker of drug resistance genes from environmental to clinically important bacteria. Current opinion in microbiology 45, 131–139 (2018)

38. Da Silva, G., Domingues, S.: Insights on the horizontal gene transfer of carbapenemase determinants in the opportunistic pathogen acinetobacter baumannii. Microorganisms 4 (3), 29 (2016)

39. Lindsay, J.A.: Staphylococcus aureus genomics and the impact of horizontal gene transfer. International Journal of Medical Microbiology 304 (2), 103–109 (2014)

40. Beck, J.M., Young, V.B., Huffnagle, G.B.: The microbiome of the lung. Translational Research 160 (4), 258–266 (2012)

41. Erb-Downward, J.R., Thompson, D.L., Han, M.K., Freeman, C.M., McCloskey, L., Schmidt, L.A., Young, V.B., Toews, G.B., Curtis, J.L., Sundaram, B., et al.: Analysis of the lung microbiome in the “healthy” smoker and in copd. PloS one 6 (2), 16384 (2011)

42. Morris, A., Beck, J.M., Schloss, P.D., Campbell, T.B., Crothers, K., Curtis, J.L., Flores, S.C., Fontenot, A.P., Ghedin, E., Huang, L., et al.: Comparison of the respiratory microbiome in healthy nonsmokers and smokers. American journal of respiratory and critical care medicine 187 (10), 1067–1075 (2013)

43. Huttenhower, C., Gevers, D., Knight, R., Abubucker, S., Badger, J.H., Chinwalla, A.T., Creasy, H.H., Earl, A.M., FitzGerald, M.G., Fulton, R.S., et al.: Structure, function and diversity of the healthy human microbiome. Nature 486 (7402), 207 (2012)

44. Li, H., Durbin, R.: Fast and accurate short read alignment with burrows–wheeler transform. Bioinformatics 25 (14), 1754–1760 (2009)

45. Jaillard, M., Tournoud, M., Meynier, F., Veyrieras, J.-B.: Optimization of alignment-based methods for taxonomic binning of metagenomics reads. Bioinformatics 32 (12), 1779–1787 (2016)

46. Jaillard, M., Lima, L., Tournoud, M., Mahé, P., van Belkum, A., Lacroix, V., Jacob, L.: A fast and agnostic method for bacterial genome-wide association studies: bridging the gap between kmers and genetic events. bioRxiv, 297754 (2018)

47. Peng, Y., Leung, H.C., Yiu, S.-M., Chin, F.Y.: Idba-ud: a de novo assembler for single-cell and metagenomic sequencing data with highly uneven depth. Bioinformatics 28 (11), 1420–1428 (2012)

48. Tarendeau, F.: Method for the specific isolation of nucleic acids of interest. US Patent WO/2014/114896 (2014)

