## Supplementary material for "Clinical metagenomics bioinformatics pipeline for the identification of hospital-acquired pneumonia pathogens antibiotic resistance genes from bronchoalveolar lavage samples"

February 14, 2020

This document presents supplementary information for the manuscript entitled "Clinical metagenomics bioinformatics pipeline for the identification of hospital-acquired pneumonia pathogens antibiotic resistance genes from bronchoalveolar lavage samples".

#### 1 Taxonomic binning: flora class composition

Figure 1 presents a Krona chart of the flora class composition. The percentages are relative to the number of genomes included in the reference database, used to learn the multi-class predictor for taxonomic read binning.

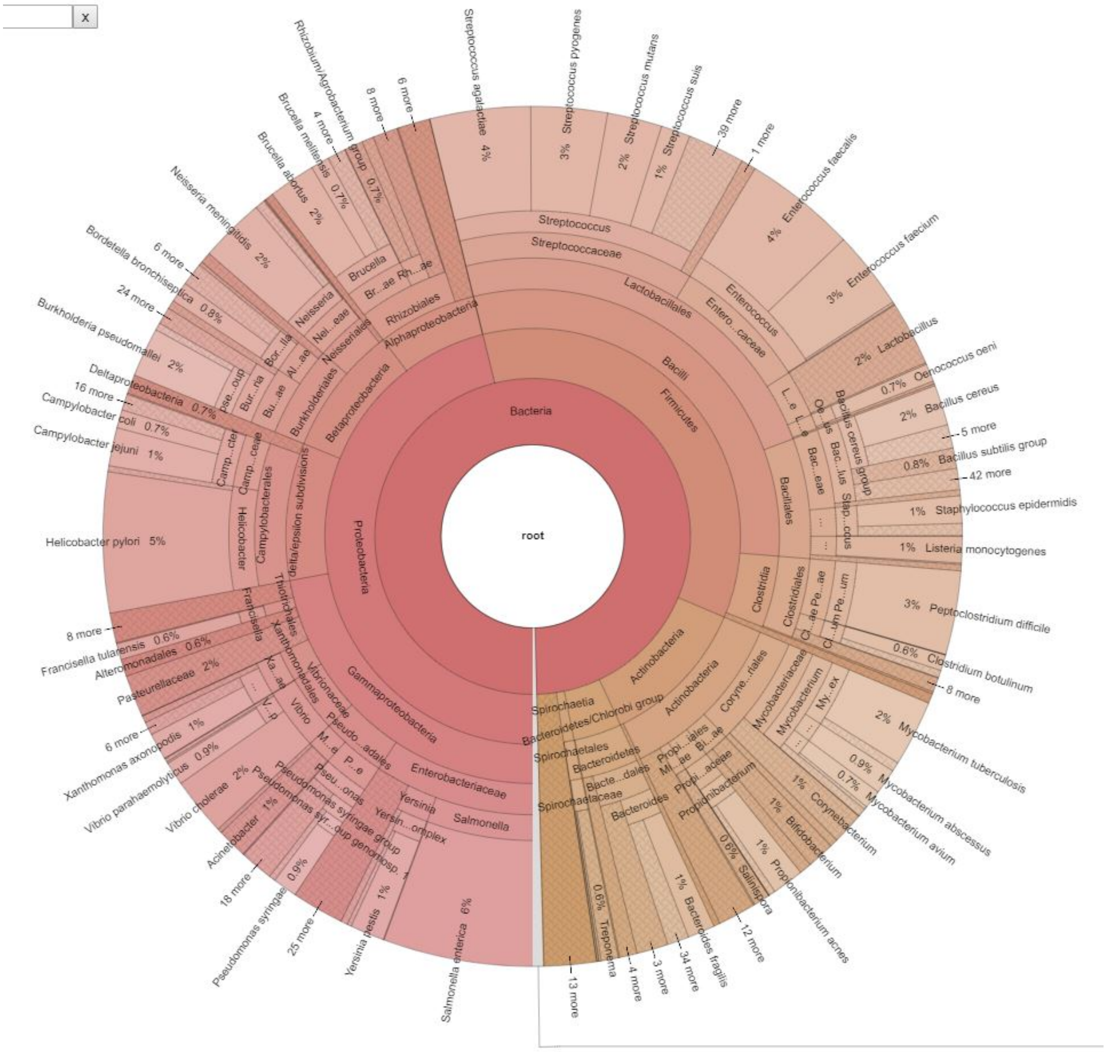


Figure 1: Composition of the flora class.

#### 2 Taxonomic binning: performance of the multi-class predictor

Figure 2 presents the confusion matrix of the classification of simulated reads from 20 strains corresponding to HAP VAP pathogens, reads simulated from genomes of the flora class, and reads simulated from the human genome. The classifier does not classify reads with uncertain predictions (“No-prediction” class). As can be seen in Figure 2, the sensitivity is above 90% for all pathogens of interest, except for *H. alvei* for which half of the reads remains unclassfied. This can be explained by the fact that we have only 17 representative genomes for this species.

#### HAP VAP taxonomic binning

| | Acinetobacter baumannii | Citrobacter freundii | Citrobacter koseri | Enterobacter aerogenes | Enterobacter cloacae | Escherichia coli | Haemophilus influenzae | Hafnia alvei | Klebsiella oxytoca | Klebsiella pneumoniae | Legionella pneumophila | Morganella morganii | Proteus mirabilis | Proteus vulgaris | Providencia stuartii | Pseudomonas aeruginosa | Serratia marcescens | Staphylococcus aureus | Stenotrophomonas maltophilia | Streptococcus pneumoniae | Flora | Human | No−prediction |
|---|---|---|---|---|---|---|---|---|---|---|---|---|---|---|---|---|---|---|---|---|---|---|---|
| Acinetobacter baumannii | 97.8 | 0 | 0 | 0 | 0 | 0 | | 0 | 0 | 0 | | 0 | | 0 | 0 | 0 | 0 | | 0 | | 1.4 | 0 | 0.7 |
| Citrobacter freundii | 0 | 96 | 0 | 0 | 0.1 | 0 | | 0 | 0 | 0 | | | | | | 0 | 0 | | 0 | | 2.4 | 0 | 1.5 |
| Citrobacter koseri | 0 | 0.1 | 90.5 | 1.6 | 0.2 | 0.2 | | 0 | 0.4 | 0.2 | | 0 | 0 | | | 0 | 0 | | 0 | | 2 | 0 | 4.9 |
| Enterobacter aerogenes | 0 | 0 | 0.1 | 94.6 | 0.2 | 0 | | 0 | 0.2 | 0.2 | | 0 | 0 | 0 | | 0 | 0 | | 0 | | 1.6 | 0 | 3.1 |
| Enterobacter cloacae | 0 | 0.4 | 0 | 0 | 90.2 | 0.1 | | 0.3 | 0.2 | 0.2 | | 0 | | 0 | 0 | 0 | 0.1 | | 0 | | 1.5 | 0 | 7 |
| Escherichia coli | | 0.1 | 0.1 | 0 | 0.3 | 94.1 | 0 | 0 | 0 | 0 | | 0 | 0.2 | | | 0 | 0 | | 0 | | 1.3 | 0 | 3.8 |
| Haemophilus influenzae | 0 | 0 | 0 | 0 | 0.1 | 0 | 96.4 | 0 | 0 | 0 | | 0 | 0 | 0 | 0 | 0 | 0 | | | | 2.4 | 0.1 | 0.9 |
| Hafnia alvei | 0 | 0.1 | 0 | 0 | 0.2 | 0.1 | | 51.5 | 0.1 | 0 | 0 | 0 | 0 | 0 | | 0 | 0 | 0 | 0 | | 6.5 | 0.2 | 41.2 |
| Klebsiella oxytoca | | 0 | 0 | 0 | 0.1 | 0 | | 0 | 96.1 | 0 | | 0 | | 0 | | 0 | 0 | | 0 | | 1.4 | 0 | 2.3 |
| Klebsiella pneumoniae | 0 | 0 | 0 | 0.1 | 0.1 | 0 | | 0 | 0.1 | 95.8 | | 0 | 0 | 0 | | 0 | 0 | | 0 | | 1.4 | 0 | 2.4 |
| Legionella pneumophila | 0 | 0 | | 0 | 0 | 0 | | 0 | 0 | 0 | 97.4 | | | 0 | | 0 | 0 | | | | 1.9 | 0.1 | 0.5 |
| Morganella morganii | 0.1 | 0 | 0 | 0 | 0.1 | 0.1 | 0 | 0 | 0 | 0 | | 94.4 | 0 | 0 | 0.2 | 0 | 0 | | 0 | | 2.1 | 0.1 | 2.9 |
| Proteus mirabilis | 0 | 0 | 0 | 0 | 0.1 | 0 | | 0 | 0 | 0 | | 0 | 92.2 | 1.1 | 0.1 | 0 | 0 | | 0 | | 2.4 | 0.1 | 3.9 |
| Proteus vulgaris | 0 | 0 | 0 | 0 | 0.1 | 0 | | 0 | 0 | 0 | 0 | 0 | 0.1 | 96.2 | 0 | 0 | 0 | 0 | | | 2.1 | 0.1 | 1.3 |
| Providencia stuartii | 0 | 0 | 0 | 0 | 0.1 | 0 | | 0 | 0 | 0 | 0 | 0 | 0.1 | 0 | 91.8 | | 0 | | | | 3 | 0.1 | 4.8 |
| Pseudomonas aeruginosa | 0 | 0 | 0 | 0 | 0 | 0 | | | 0 | 0 | | 0 | | | | 95 | 0 | | 0.2 | | 2.4 | 0 | 2.2 |
| Serratia marcescens | | 0 | 0 | 0 | 0.1 | 0 | 0 | 0 | 0 | 0 | | 0 | 0 | 0 | | 0 | 92.6 | | 0 | | 2.6 | 0 | 4.5 |
| Staphylococcus aureus | 0 | 0 | | 0 | 0 | 0 | | 0 | 0 | 0 | 0 | | | | | | 0 | 96.9 | | 0 | 2.2 | 0.1 | 0.6 |
| Stenotrophomonas maltophilia | 0 | 0 | 0 | 0 | 0.1 | 0 | | | 0 | 0 | | | 0 | | | 0.1 | 0 | | 90.8 | | 4.1 | 0 | 4.7 |
| Streptococcus pneumoniae | 0 | 0 | 0 | 0 | 0 | 0 | | 0 | 0 | 0 | | | | | | 0 | 0 | | | 97.3 | 2 | 0.1 | 0.5 |
| Flora | 0 | 0 | 0 | 0 | 0.1 | 0.3 | 0 | 0 | 0 | 0 | 0 | 0 | 0 | 0 | 0 | 0.4 | 0 | 0.1 | 0.2 | 0.6 | 43.6 | 0.7 | 53.7 |
| Human | 0 | 0 | 0 | 0 | 0 | 0 | | 0 | 0 | 0 | 0 | 0 | 0 | 0 | 0 | 0 | 0 | 0 | 0 | 0 | 2.5 | 92.9 | 4.5 |

Figure 2: Multi-class predictor for taxonomic binning: confusion matrix. In row are the reference labels (labels of the simulated strains) and in columns the predicted labels. “No-prediction” means that the predictor did not classidied the read pair.

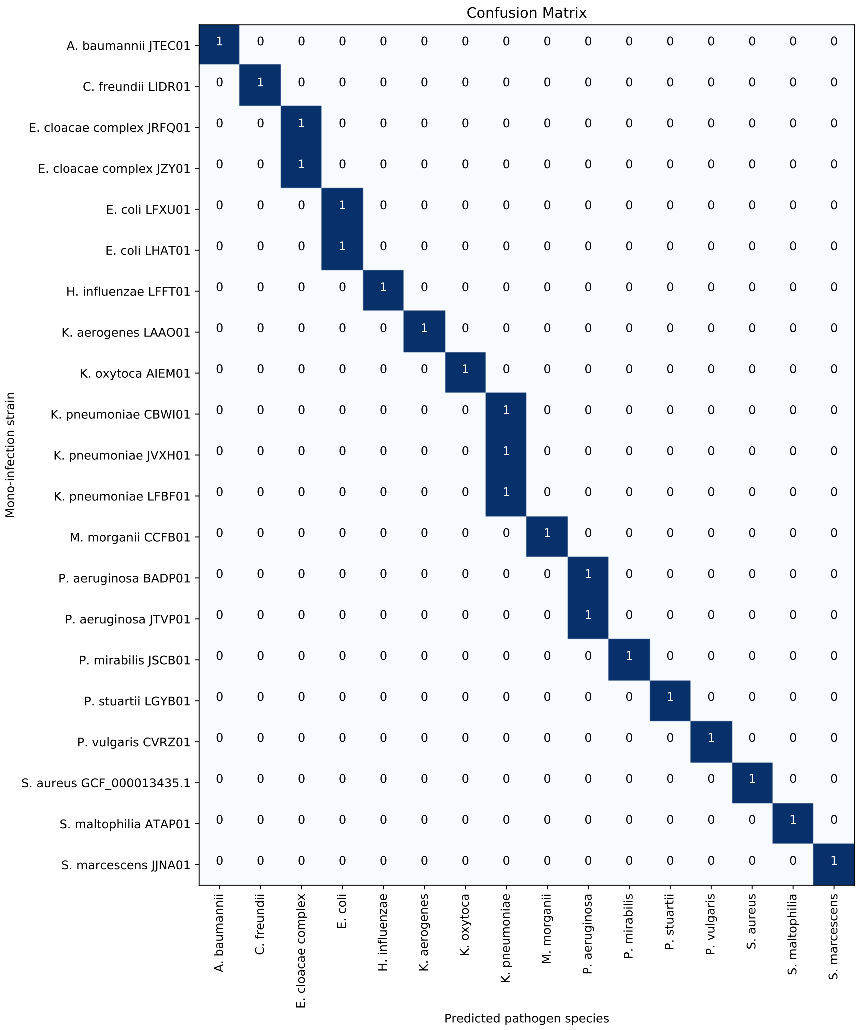


Figure 3: Monomicrobial simulations confusion matrix for pipeline TBo and pipeline TBwDM (same performance).

### 3 Monomicrobial and polymicrobial infection simulations: species and ARG detection results

Figure 3 and 4 present the confusion matrices of our bioinformatics pipelines species detection step, respectively in the monomicrobial and in the polymicrobial simulations. Both pipelines achieve the same performance.

Figure 5 and 6 illustrate the marker recovery and assignment performance of our bioinformatics pipelines, in the case of simulated polymicrobial infections. Both figures provide ARG detection details for each pathogen in each scenario. At first, we can visualize that ARG detection recall is generally improved by pipeline TBwDM vs. pipeline TBo. Then, pathogen coverage configurations, which can be balanced (see

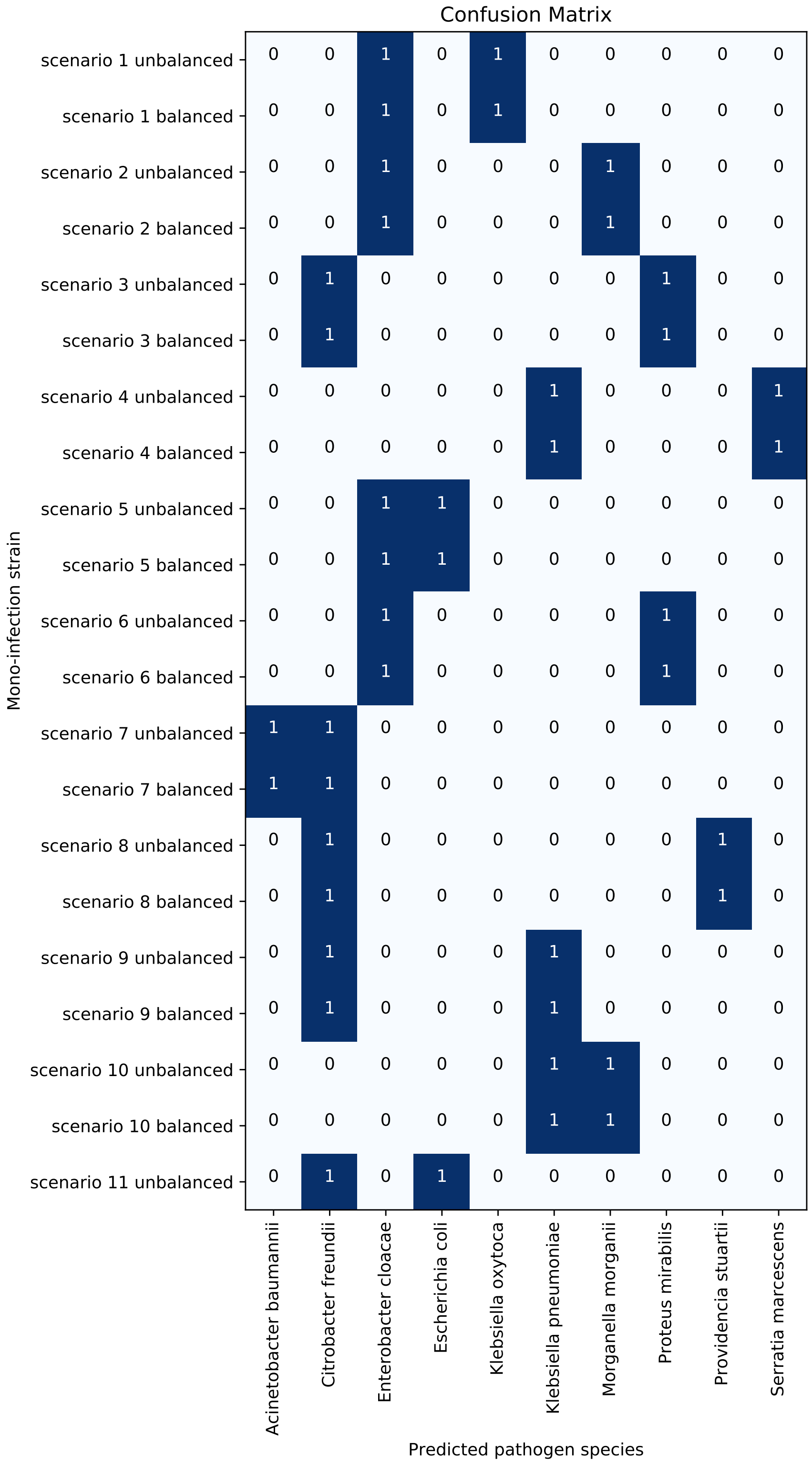


Figure 4: Polymicrobial simulations confusion matrix for pipeline TBo and pipeline TBwDM (same performance). *E. cloacae* complex is not detected in scenario 11 unbalanced, leading to a false negative since it was included in the 3-pathogens scenario 11 as the lowest concentrated pathogen (coverage 0.45X).

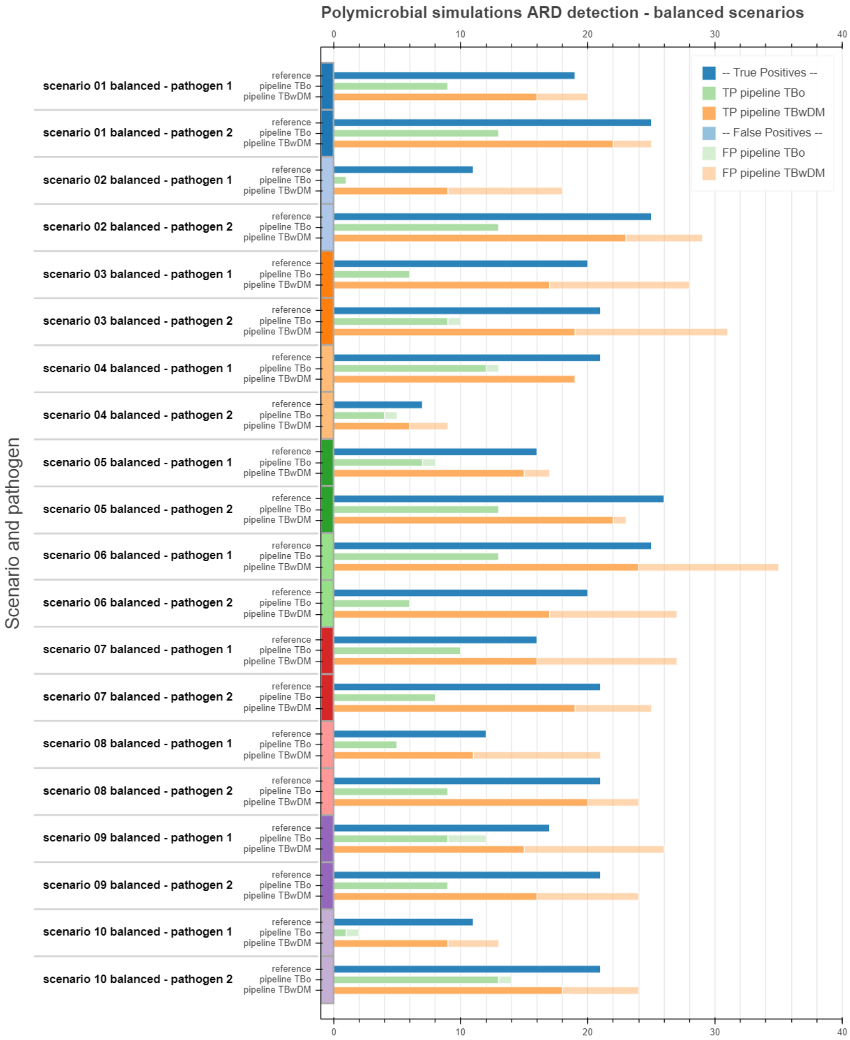


Figure 5: Balanced polymicrobial simulations ARG detection results for pipeline TBo and pipeline TBwDM. Reference lines state for the true number of ARG in each scenario / pathogen. Pipelines TBo and TBwDM lines show ARG detection results, with solid color standing for true positives, and faded color standing for false positives. False negatives can be deduced by subtracting true positives from the reference.

Figure 5) or unbalanced (see Figure 6), lead to different results, the latter tending to improve precision on the higher-coverd pathogen (pathogen 2), while decreasing the lower-covered pathogen (pathogen 1) recall at the same time. Table 1 provide those precision and recall values, for each 2 pathogens polymicrobial scenario.

To better evaluate how much ARG assignment can impact the performance, we also analyzed the ARG detection without pathogen assignment step, ARG being considered at the sample level and not at the pathogen level. Figure 7 illustrate the ARG detection results for each each polymicrobial scenario. While pipeline TBo performances remain quite stable, pipeline TBwDM precision and recall increase when we look at

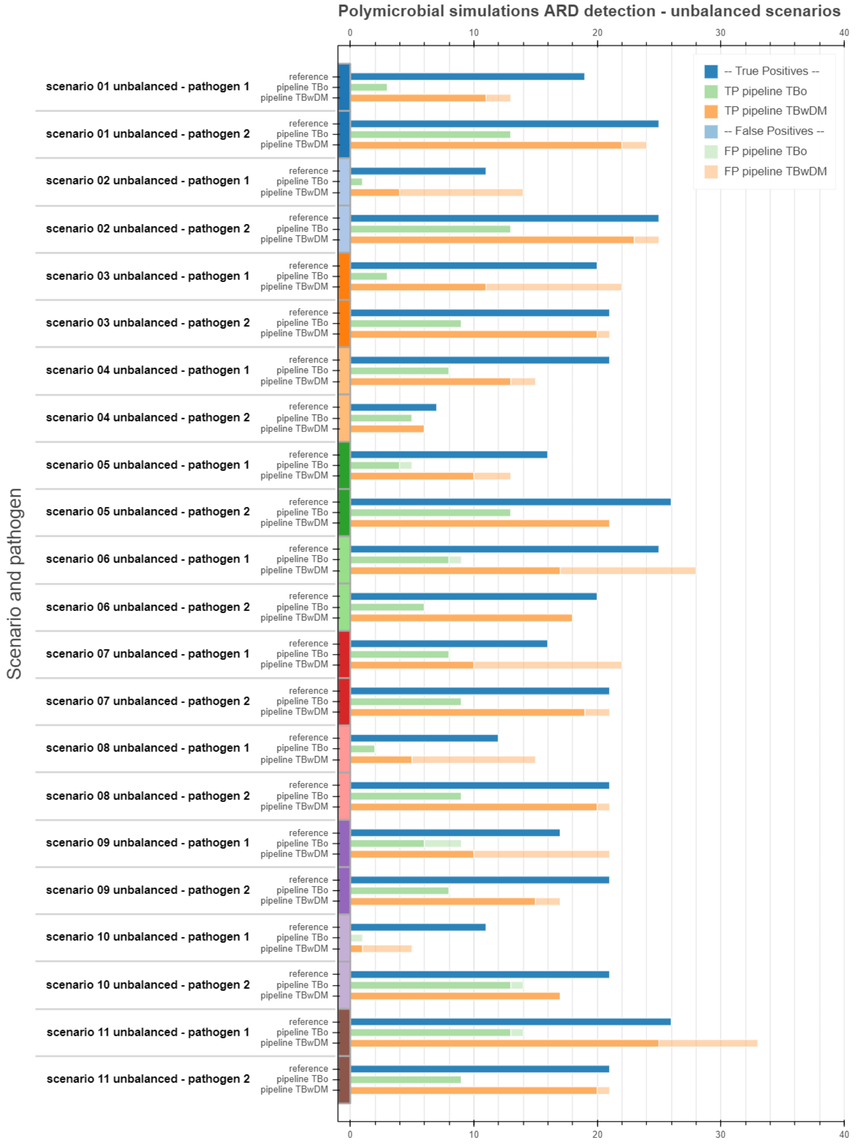


Figure 6: Unbalanced polymicrobial simulations ARG detection results for pipelines TBo and TBwDM. Reference lines state for the true number of ARG in each scenario / pathogen. Pipelines TBo and TBwDM lines show ARG detection results, with solid color standing for true positives, and faded color standing for false positives. False negatives can be deduced by subtracting true positives from the reference.

the sample level.

Finally, we use MetaCherchant to provide some valuable insight on the genomic context of the ARG which are detected by our bioinformatics pipeline TBwDM. What we suggest is to focus on these detected ARG, and to label their subgraphs by blasting the related unitigs against a subset of our pneumonia RDB, by restricting it to the detected pathogens. As an example, Figure 8 to 10 illustrate different genomic contexts that we met in our simulations, including an example of false assignment.

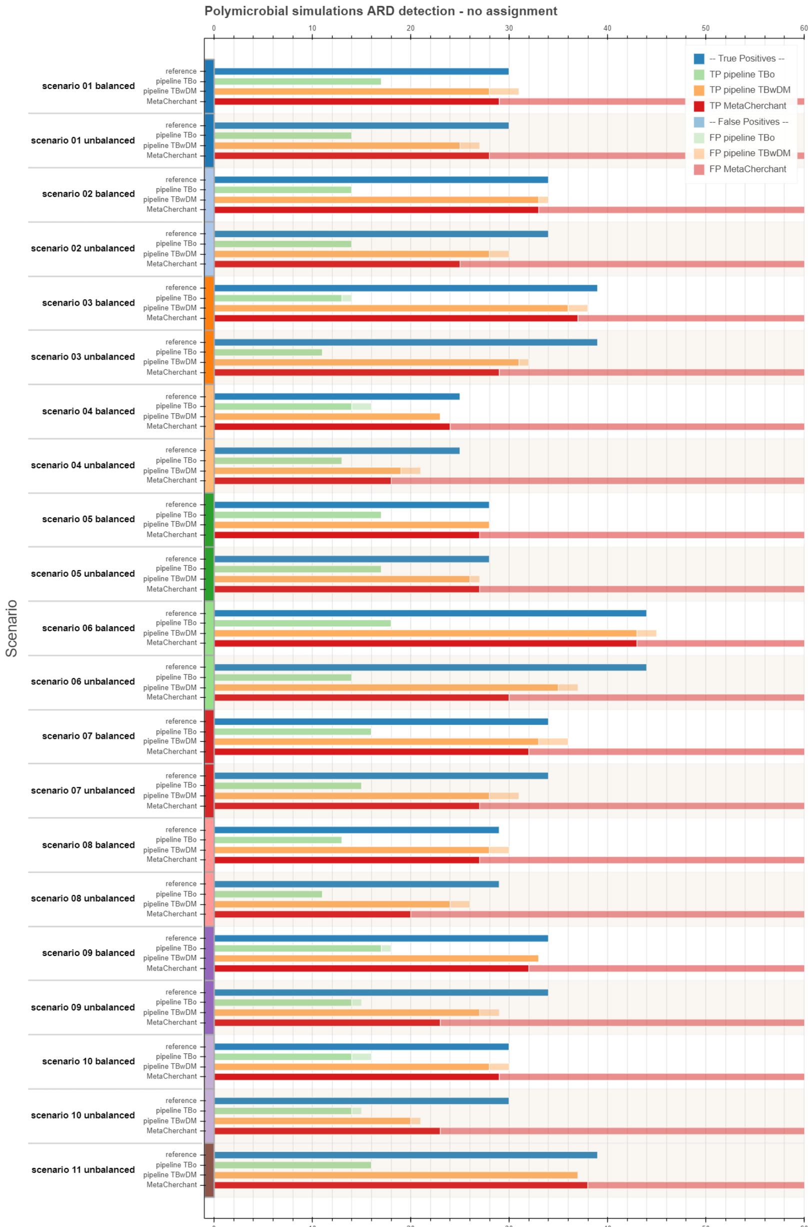


Figure 7: Polymicrobial simulations ARG detection results, when ARG are considered at the sample level, i.e. not assigned to some pathogens. Reference lines state for the true number of ARG in each sample (all pathogens mixed). Pipelines TBo, TBwDM and MetaCherchant lines show ARG detection results, with solid color standing for true positives, and faded color standing for false positives. Exact total number of MetaCherchant false positives is not provided, because we did not implement any output graphs post-processing to select one unique best-hit-variant ARG per ARG name, and so we believe that it would not be fair. False negatives can be deduced by subtracting true positives from the reference.

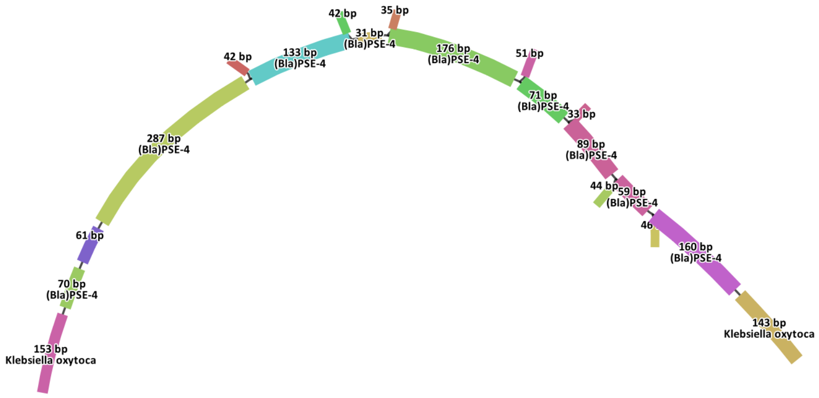


Figure 8: Polymicrobial simulations ARG genomic context understanding using MetaCherchant. Subgraphs are drawn in the neighbourhood of ARG that are detected by our pipeline TBwDM, and related unitigs are blasted against a subset of our pneumonia RDB based on the detected pathogens. Scenario 1 unbalanced, ARG *(Bla)PSE-4* is correctly assigned to *K. oxytoca*.

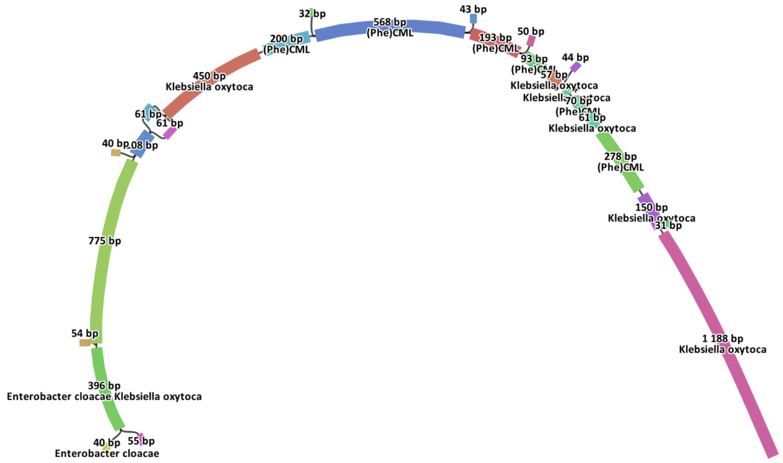


Figure 9: Polymicrobial simulations ARG genomic context understanding using MetaCherchant. Subgraphs are drawn in the neighbourhood of ARG that are detected by our pipeline TBwDM, and related unitigs are blasted against a subset of our pneumonia RDB based on the detected pathogens. Scenario 1 unbalanced, ARG *(Phe)CML* is correctly assigned to both pathogens pathogens *E. cloacae complex* and *K. oxytoca*, even though *E. cloacae complex* is further from the ARG position.

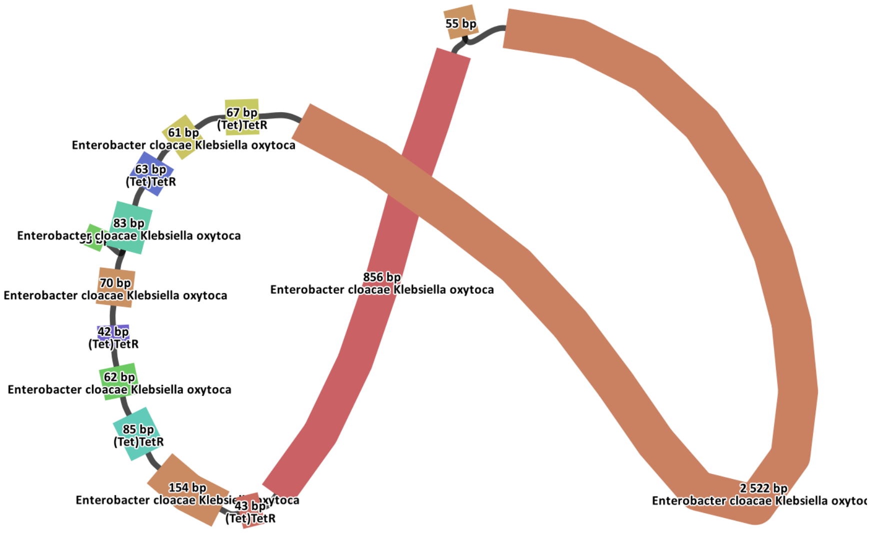


Figure 10: Polymicrobial simulations ARG genomic context understanding using MetaCherchant. Subgraphs are drawn in the neighbourhood of ARG that are detected by our pipeline TBwDM, and related unitigs are blasted against a subset of our pneumonia RDB based on the detected pathogens. Scenario 1 unbalanced, ARG *(Tet)TetR* is correctly assigned to *E. cloacae complex* by our pipeline TBwDM, but MetaCherchant also finds *K. oxytoca* in the genomic neighbourhood of the ARG.

| | | | Pipeline TBo | | Pipeline TBwDM | |
|---|---|---|---|---|---|---|
| Scenario | Path. # | # ARG | Prec. | Rec. | Prec. | Rec. |
| Scen. 1 balanced | 1 | 19 | 100 | 47.4 | 80 | 84.2 |
| Scen. 1 balanced | 2 | 25 | 100 | 52 | 88 | 88 |
| Scen. 2 balanced | 1 | 11 | 100 | 9.1 | 50 | 81.8 |
| Scen. 2 balanced | 2 | 25 | 100 | 52 | 79.3 | 92 |
| Scen. 3 balanced | 1 | 20 | 100 | 30 | 60.7 | 85 |
| Scen. 3 balanced | 2 | 21 | 90 | 42.9 | 61.3 | 90.5 |
| Scen. 4 balanced | 1 | 21 | 92.3 | 57.1 | 100 | 90.5 |
| Scen. 4 balanced | 2 | 7 | 80 | 57.1 | 66.7 | 85.7 |
| Scen. 5 balanced | 1 | 16 | 87.5 | 43.8 | 88.2 | 93.8 |
| Scen. 5 balanced | 2 | 26 | 100 | 50 | 95.7 | 84.6 |
| Scen. 6 balanced | 1 | 25 | 100 | 52 | 68.6 | 96 |
| Scen. 6 balanced | 2 | 20 | 100 | 30 | 63.0 | 85 |
| Scen. 7 balanced | 1 | 16 | 100 | 62.5 | 59.3 | 100 |
| Scen. 7 balanced | 2 | 21 | 100 | 38.1 | 76 | 90.5 |
| Scen. 8 balanced | 1 | 12 | 100 | 41.7 | 52.4 | 91.7 |
| Scen. 8 balanced | 2 | 21 | 100 | 42.9 | 83.3 | 95.2 |
| Scen. 9 balanced | 1 | 17 | 75 | 52.9 | 57.7 | 88.2 |
| Scen. 9 balanced | 2 | 21 | 100 | 42.9 | 66.7 | 76.2 |
| Scen. 10 balanced | 1 | 11 | 50 | 9.1 | 69.2 | 81.8 |
| Scen. 10 balanced | 2 | 21 | 14 | 92.9 | 75 | 85.7 |
| Scen. 1 unbalanced | 1 | 19 | 100 | 15.8 | 84.6 | 57.9 |
| Scen. 1 unbalanced | 2 | 25 | 100 | 52 | 91.7 | 88 |
| Scen. 2 unbalanced | 1 | 11 | 100 | 9.1 | 28.6 | 36.4 |
| Scen. 2 unbalanced | 2 | 25 | 100 | 52 | 92 | 92 |
| Scen. 3 unbalanced | 1 | 20 | 100 | 15 | 50 | 55 |
| Scen. 3 unbalanced | 2 | 21 | 100 | 42.9 | 95.2 | 95.2 |
| Scen. 4 unbalanced | 1 | 21 | 100 | 38.1 | 86.7 | 61.9 |
| Scen. 4 unbalanced | 2 | 7 | 100 | 71.4 | 100 | 85.7 |
| Scen. 5 unbalanced | 1 | 16 | 80 | 25 | 76.9 | 62.5 |
| Scen. 5 unbalanced | 2 | 26 | 100 | 50 | 100 | 80.8 |
| Scen. 6 unbalanced | 1 | 25 | 88.9 | 32 | 60.7 | 68 |
| Scen. 6 unbalanced | 2 | 20 | 100 | 30 | 100 | 90 |
| Scen. 7 unbalanced | 1 | 16 | 100 | 50 | 45.5 | 62.5 |
| Scen. 7 unbalanced | 2 | 21 | 100 | 42.9 | 90.5 | 90.5 |
| Scen. 8 unbalanced | 1 | 12 | 100 | 16.7 | 33.3 | 41.7 |
| Scen. 8 unbalanced | 2 | 21 | 100 | 42.9 | 95.2 | 95.2 |
| Scen. 9 unbalanced | 1 | 17 | 66.7 | 35.3 | 47.6 | 58.8 |
| Scen. 9 unbalanced | 2 | 21 | 100 | 38.1 | 88.2 | 71.4 |
| Scen. 10 unbalanced | 1 | 11 | 0 | 0 | 20 | 9.1 |
| Scen. 10 unbalanced | 2 | 21 | 92.9 | 61.9 | 100 | 81.0 |

Table 1: ARG detection performance detail for 2 pathogens polymicrobial simulations, i.e. scenarios 1 to 10.

### 4 Real data: phenotypic antibiotic susceptibility results

Samples 1 and 2 are *E. coli* infections while samples 3 and 4 are polymicrobial infections: *K. pneumoniae* and *H. influenzae* (sample 3) and *E. coli* and *K. aerogenes* (sample 4). Figure 11 presents a Venn Diagram with detected ARG on BAL samples with TBo (blue circle), TBwDM (yellow circle), and on isolate sequences (green circle).

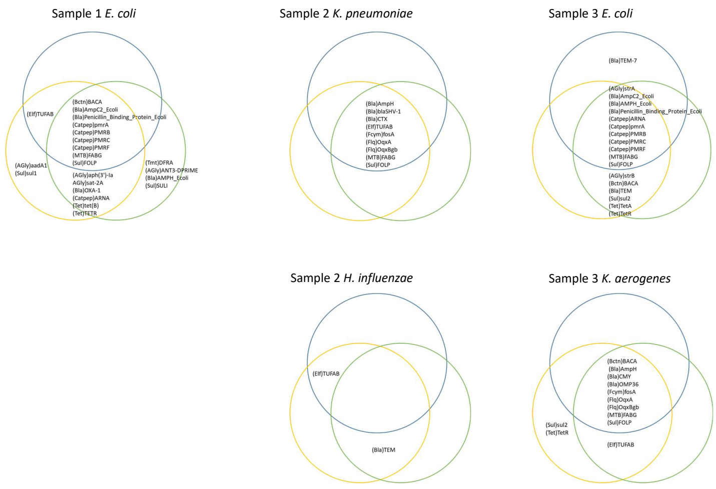


Figure 11: **ARG markers detection.** Venn Diagram with detected ARG markers on BAL samples with TBo (blue circle), TBwDM (yellow circle), and on isolate sequences (green circle). Samples are presented in column, 1 row per strain for polymicrobial samples.
